# Accelerating Antibody Design with Active Learning

**DOI:** 10.1101/2022.09.12.507690

**Authors:** Seung-woo Seo, Min Woo Kwak, Eunji Kang, Chaeun Kim, Eunyoung Park, Tae Hyun Kang, Jinhan Kim

**Author notes:** Equal contribution (Seung-woo Seo, Min Woo Kwak). Work performed while at Standigm (Seung-woo Seo).

## Abstract

Discovering therapeutic antibody starts by screening antibody library of phage-displayed, transgenic mouse or human B cells. The coverage of those kinds of libraries in the entire antibody sequence space is small; thus, the result highly depends on the quality of the library. Exploring sequence space by mutating a template antibody is also impossible to even with the state-of-the-art screening methods because of the cost. Deep learning helped with its pattern recognition nature to predict target binding, which is only applied to HCDR3 because the number of data deep learning needs increases exponentially. We construct a sequence generation model with transfer learning and active learning to leverage deep learning even in data deficiency. With only six thousands data, the generative model finds nine binding antibody sequences at least per antigen with novel HCDR3.

## 1 Introduction

Therapeutic antibodies have been the pharmaceutical industry’s focus since 1986, when the FDA approved the first monoclonal antibody (mAb) product. Four therapeutic antibodies are included in the top-10 bestselling drugs in 2021 [1], and the number of FDA-approved therapeutic antibodies is more than a hundred at present[2]. mAb has two chains (heavy and light) and two regions (variable and constant) for each chain. Variable heavy (VH) and variable light (VL) chains are regarded as mainly responsible for target specificity because the variable regions are enormously diverse among antibodies for adapting to the target molecule(antigen). Due to this fact, most screening efforts are focused on variable regions in terms of target specificity, which consist of approximately 220 amino acids comprising 114 amino acids for VH and 110 amino acids for VL[3]. Even with the most powerful high-throughput screening method, it is impossible to explore all possible sequences of a variable region, which can be calculated by *A^L^* where *A* is the number of amino acids, 20, and *L* denotes the length of protein: 220 for antibody. This number would significantly decrease when considering site-specific amino acid frequency in constant regions. However, considering generation or optimization of heavy chain complementarity-determining regions 3 (HCDR3) only, which has fifteen amino acids long in average, must explore 10^8^ sequences. Nevertheless, a deep mutational scanning-based mutagenesis library is only reachable to 10^4^ sequences [4].

Because finding a novel and effective therapeutic antibody in the vast sequence space is costly and laborious, deep learning has been considered one of the powerful tools [5]. Currently, deep learning shows significant progress in many fields such as image, natural language [6], and protein structure [7, 8]. In addition, the recent studies [4, 9, 10] show deep learning is extended to drug discovery such as finding a binding between HCDR3 and target antigen. For example, the deep learning model is applied to screen *in-scilico* optimized HCDR3 sequences from library. The screening library by introducing site-directed mutagenesis were generated by mutating every single site of HCDR3. However, the limitation of preceding works is that those were concentrated only the selection of optimized HCDR3 depending on a given template sequence. because the number of data that the deep learning model needs will increase exponentially as more extended regions of interest, such as whole variable regions. It is called *the curse of dimensionality* in the machine learning theory [6].

In this paper, we construct the therapeutic antibody sequence design method, which is library-free, actively exploring the sequence space with the orchestration of deep learning and biological assay. To design the whole sequence of the variable region, we use the generative model rather than a supervised model, which only predicts target-specific or not. The generative model can design novel therapeutic antibody sequences by estimating the other antibodies’ features. As mentioned above, generating a whole sequence of the variable region would need a colossal amount of data because of *the curse of the dimensionality*. To cope with the lack of data issue, we apply the transfer learning method [11], which pretrains the model with the large dataset to learn general representations of antibodies. Then fine-tune to target-specific data. After training the deep learning model and sampling new sequences from the model, ELISA experimentally validated the generated antibodies through antigen-antibody binding. Furthermore, the model training, generation, and biological validation make a feedback loop by giving the assay results to the model for training (Fig. 1). This loop of a train, querying, and validation is called the *active learning* [12] which is effective when the labeled data is scarce. As a result, active learning effectively finds antibody sequences with unique HCDR3 sequences and the other regions. The number of discovered sequences is increased while the further iteration is done.

**Figure 1:**
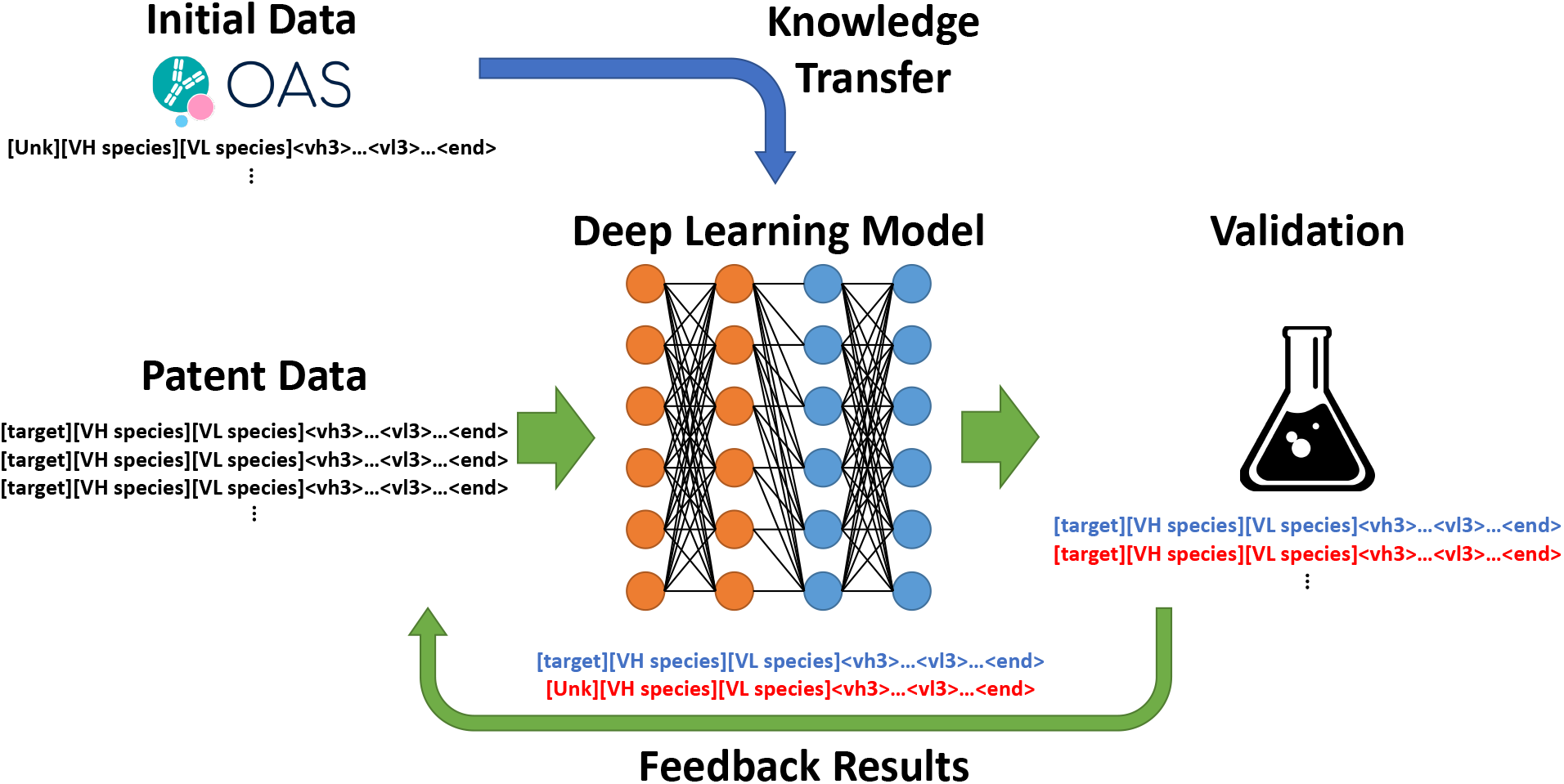
Deep learning train and generation of antibody sequences, ELISA validation and retrain loop

In brief, the contributions of our work are three folds. We construct the data-efficient framework that can discover novel therapeutic antibodies by combining deep learning models and biological lab experiments. On top of the suggested framework, our therapeutic antibody design becomes limitless over HCDR3 with a deep learning model.

## 2 Results

### 2.1 Transfer general feature of antibodies from pretraining to finetuning

The amino acid sequence of VH-VL region is recombined from V(D)J gene segment in activated B-cells [3]. The V(D)J recombination means that VH and VL have three and two independent segments respectively. According to previous deep learning researches on proteins [8, 13, 14, 15, 16, 17], the existence of independent regions in a protein is negligible. Therefore, we assume that there are sequential dependencies from each amino acid to its previous one in a whole antibody sequence. To approximate the dependencies using the deep learning model, we uses GPT-2 architecture for natural language modelling [18]. GPT-2 is the state-of-the-art deep learning architecture in language generation. Basically, GPT-2 predicts the probability of next amino acid from the given sequence. By running GPT-2 recurrently predicting and choosing amino acid with the predicted probability, we can generate the antibody sequences.

GPT-2 has millions of parameters which is far larger than our patent dataset. As a result, pretrained GPT-2 learned general features of natural antibodies such as the distribution of length of HCDR3[19, 20] and theoretical isoelectric point (pI) value. As shown in 2, HCDR3 length and calculated pI values[21, 22, 23] of generated sequences from pretrained GPT2 are comparable to those of trained sequences from OAS dataset. It is worth noting that there were no constraints about regions or physicochemical properties. Our result shows that the higher-level properties like the length of HCDR3 and pI values can be recovered from the sequential dependency of amino acid.

**Figure 2:**
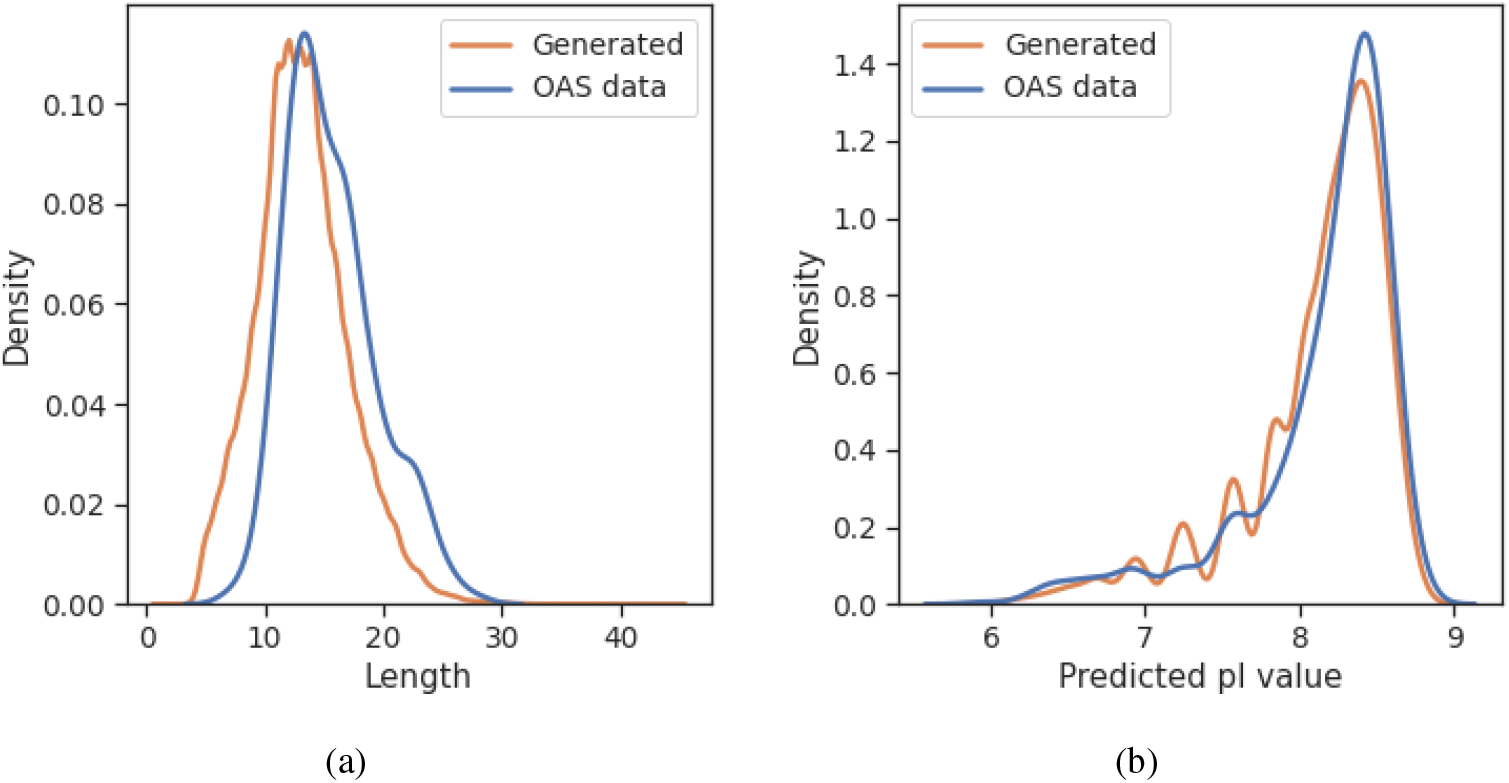
Distribution of (a) HCDR3 length and (b) predicted pI of generated and OAS test data. The distributions are smoothed by Gaussian kernel density estimation.

OAS dataset has no binding antigen data thus, generated sequences from pretrained model are not intended to have target specificity. In order to meet the intended purpose, other set of data are required such as a pairing data of antigen and antibodies. Due to lack of public data, approximately 6000 antibodies including antibody with its target antigen are curated from the patents which covered 15 oncology targets. Each target has variety ranges in number of antibody sequences. It varied from few to 1000 antibodies per each target. For example, programmed cell death protein 1(PD-1) has 1313 antibodies corresponding to PD-1 protein. On the other hand, the fewest number of antibody sequences were for MET(Mesenchymal Epithelial Transition) which only has less than 10 antibodies. To learn target specificity, model is trained on target-specified dataset with pretrained model weights. But training on the patent dataset does not guarantee that pretrained knowledge is transferred to finetuned model. Thus, we compare learning curves of training from-scratch with pretrained model. As shown in figure 3, validation loss on patent dataset converge faster and lower at finetuned model than from-scratch model. In conclusion, transfer learning accelerates more than four times in training speed until converge and give model with more performance.

**Figure 3:**
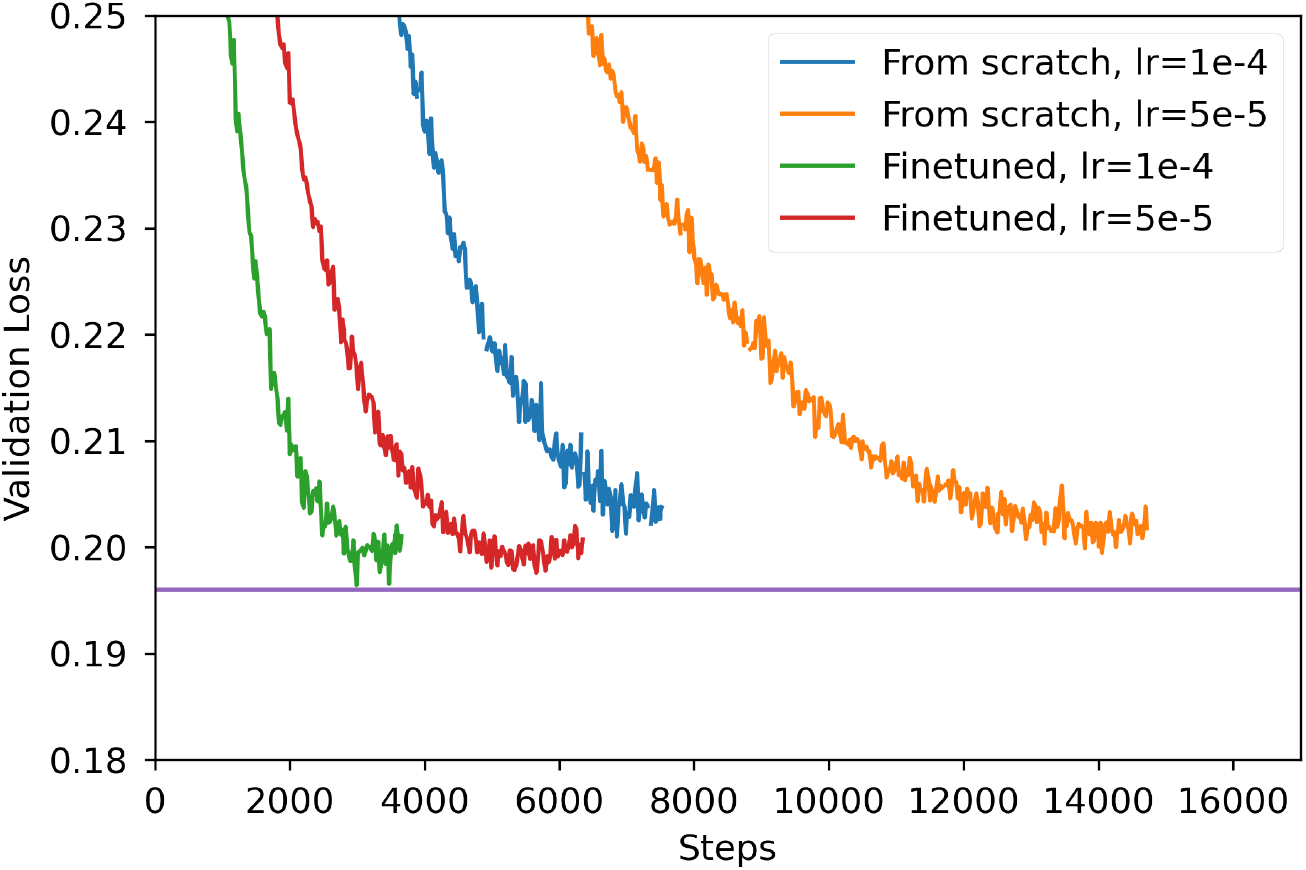
Validation loss curve during two different start condition of training. One is start from the scratch, and the other is pretrained on OAS dataset. Finetuing model shows faster convergence to find its minima and reaches to lower validation loss than train from scratch.

### 2.2 Generate target- and species-specific antibodies

Sequence generation using deep learning method has many suggested algorithms such as guided decoding [24], trainable decoding [25], gradient-based prompt design [26], and heuristic prompts [27].

In this work, we apply the Conditional Transformer(CTRL) algorithm suggested by Keskar et al. [28]. CTRL is started from a pretrained model and finetune the model using a control tag. CTRL can be seamlessly applied to our OAS pretrained model by adding target names at the front of sequences. Furthermore, we control GPT-2 to generate a human frame region by tagging species of sequences. We classified sequence species by using ANARCI [29] with IMGT. Proof on the effect of CTRL on target binding is unfeasible without accurate binding predictions. Alternatively, OAS pretrained model used to generate sequences in three conditions, without a tag, with human, or mouse tag to validate the effect of CTRL method. Without a control tag, the ratio of human generated sequences is almost 0 at VH and VL frames. However, more than a half of CTRL applied GPT-2 generated sequences are classified as human frame in both VH-VL regions in table 1. By simple changing, human to mouse tag, the model generates different frames. With mouse tag, CTRL gives more ratio than human case. This is because OAS have mouse sequences twice more than human. Among the generated sequences, a hundred sequences with the lowest loss are refined not to have duplicates in training, validation and test dataset, and passed to experimental validation.

**Table 1:** Effect of CTRL method in control generated frame species. Numbers are the ratio of target species to total generated sequences from model.

| Tag | VH |  | VL |  | VH-VL |  |
| --- | --- | --- | --- | --- | --- | --- |
|  | Human | Mouse | Human | Mouse | Human | Mouse |
| No tag | 0.015 | 0.402 | 0.076 | 0.800 | 0.004 | 0.340 |
| Human | 0.607 |  | 0.937 |  | 0.574 |  |
| Mouse. |  | 0.667 |  | 0.987 |  | 0.656 |

### 2.3 Experimental validation of the generative model

To validate the performance of generative model, ML model based on CTRL applied GPT2 was utilized to generate target specific antibodies. Then, generated antibody sequences were investigated their binding activity followed by serials of experimental testing. For this purpose, programmed cell death protein 1(PD-1) and programmed death-ligand 1(PD-L1) were selected which are the promising immuno-oncology targets and already have several approved therapeutic antibodies and still under active clinical investigations.

During three rounds of sequence generation, experimental validation and feedback to data loop, 218 clones of AI-designed anti-PD-1, and 183 clones of AI-designed anti-PD-L1 were screened to figure out whether those are antigen binders or not. First, bacterial expression system was used to verify designed antibodies because it requires less effort, time, and money in a high-throughput manner, compared with mammalian expression system. OmpA (Outer membrane protein A) signal peptide let scFv proteins be secreted to bacteria periplasm and the bacterial outer membrane and the peptidoglycan layer were removed to obtain the periplasm. ScFv proteins were fused to HA tag at C-terminal, and therefore soluble scFv proteins in periplasmic fraction bound to target antigen could be detected with HRP-conjugated anti-HA antibody by ELISA assay. By reading 450 nm absorbance with microplate reader, antibodies showing absorbance of over 0.1 was regarded as target antigen binders. Consequently, nine anti-PD-1 antibodies and ten anti-PD-L1 antibodies showed binding to the corresponding antigens in initial screening. Simultaneously, dot blot was performed using the same periplasmic fraction to confirm periplasmic expression of each candidate.

Furthermore, purified positive candidates were further evaluated in dose dependent manner in an ELISA assay. Nivolumab (Opdivo®) and pembrolizumab (Keytruda®) were used as anti-PD-1 positive controls, and Atezolizumab, Avelumab, and Durvalumab were used as anti-PD-L1 positive controls. In-house generated non-specific (anti-LPA2) antibody was used as a negative control. Purified antibodies including positive and negative controls were serially diluted from 1000 nM in duplicate. In a large scale, two anti-PD-L1 antibodies (clone 162 and 163) were failed to achieve by developed protocol described in material and method section 3. The purified 17 antibodies were further investigated their binding activities to its corresponding antigen in dose dependent manner. The data including purification and binding active were summarised in table 2. Five antibodies of them did not display binding to target antigens, four in anti-PD-1 and one in anti-PD-L1, respectively. Consequently, 14 AI-designed antibodies were turned out to be positive clones which bind to the target antigens. Anti-PD-1 antibody clone 77 (0.094 nM) showed lower EC50 (nM) value than anti-PD-1 positive control, which are nivolumab (0.93 nM) and pembrolizumab (4.37 nM). There was no higher binder than the PD-L1 control antibodies, atezolizumab (0.17 nM), avelumab (0.35 nM), and durvalumab (0.71 nM) but the anti-PD-L1 antibody clone 187 (0.82 nM) showed similar binding activity to PD-L1 as durvalumab. In summary, the current study involving AI-based de novo generation and in vitro experimental study successfully identified novel antibodies against PD-1 and PD-L1.

**Table 2:**
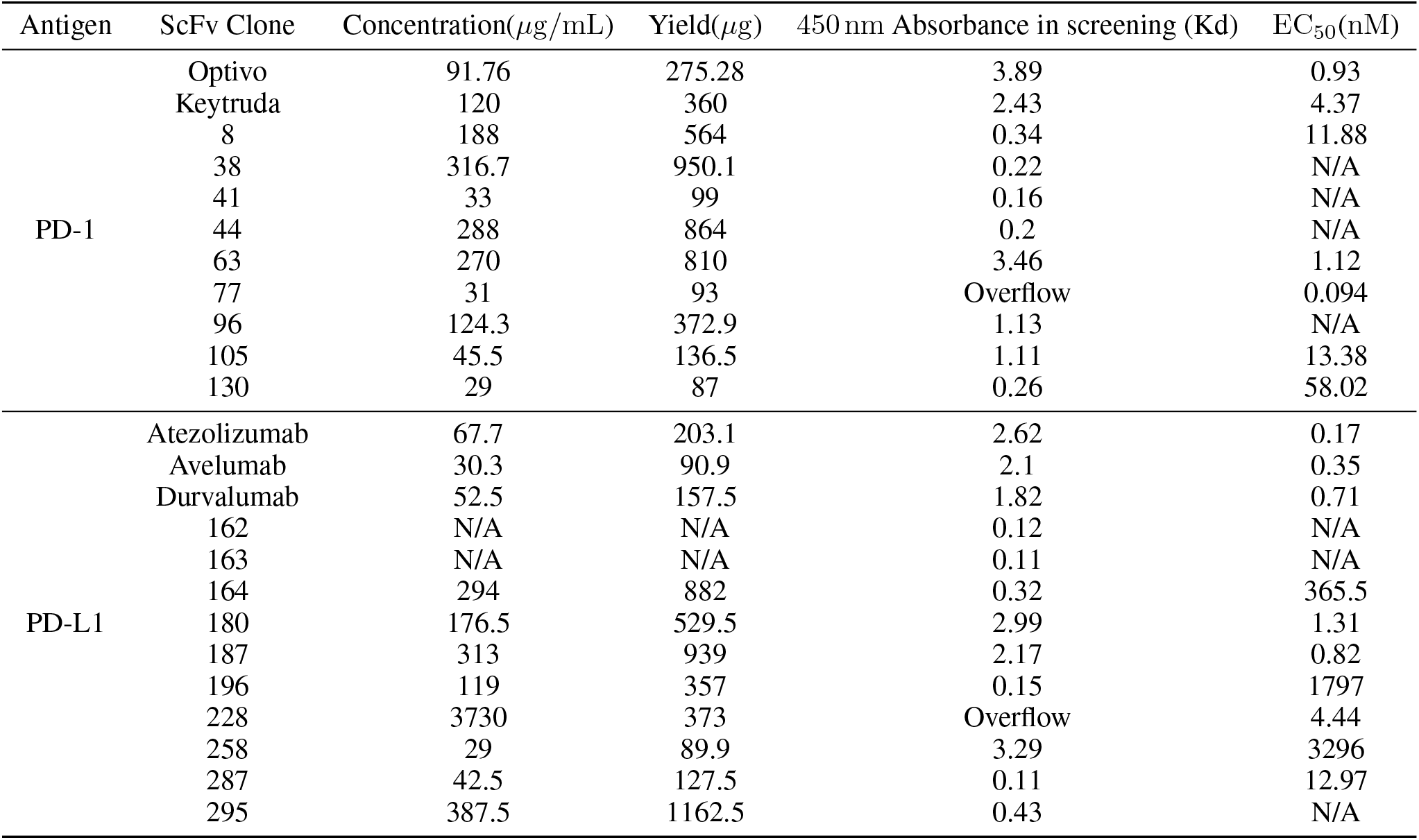
Concentration, yield, measured Kd value after purification and absorbance in screening of individual scFv clones. N/A means not analyzed.

| Antigen | ScFv Clone | Concentration($\mu\text{g/mL}$) | Yield($\mu\text{g}$) | 450 nm Absorbance in screening (Kd) | EC<sub>50</sub>(nM) |
| --- | --- | --- | --- | --- | --- |
| PD-1 | Optivo | 91.76 | 275.28 | 3.89 | 0.93 |
|  | Keytruda | 120 | 360 | 2.43 | 4.37 |
|  | 8 | 188 | 564 | 0.34 | 11.88 |
|  | 38 | 316.7 | 950.1 | 0.22 | N/A |
|  | 41 | 33 | 99 | 0.16 | N/A |
|  | 44 | 288 | 864 | 0.2 | N/A |
|  | 63 | 270 | 810 | 3.46 | 1.12 |
|  | 77 | 31 | 93 | Overflow | 0.094 |
|  | 96 | 124.3 | 372.9 | 1.13 | N/A |
|  | 105 | 45.5 | 136.5 | 1.11 | 13.38 |
|  | 130 | 29 | 87 | 0.26 | 58.02 |
| PD-L1 | Atezolizumab | 67.7 | 203.1 | 2.62 | 0.17 |
|  | Avelumab | 30.3 | 90.9 | 2.1 | 0.35 |
|  | Durvalumab | 52.5 | 157.5 | 1.82 | 0.71 |
|  | 162 | N/A | N/A | 0.12 | N/A |
|  | 163 | N/A | N/A | 0.11 | N/A |
|  | 164 | 294 | 882 | 0.32 | 365.5 |
|  | 180 | 176.5 | 529.5 | 2.99 | 1.31 |
|  | 187 | 313 | 939 | 2.17 | 0.82 |
|  | 196 | 119 | 357 | 0.15 | 1797 |
|  | 228 | 3730 | 373 | Overflow | 4.44 |
|  | 258 | 29 | 89.9 | 3.29 | 3296 |
|  | 287 | 42.5 | 127.5 | 0.11 | 12.97 |
|  | 295 | 387.5 | 1162.5 | 0.43 | N/A |

## 3 Method and Materials

### 3.1 Dataset

Observed antibody space(OAS) is cleansed and IMGT numbering applied dataset containing VH and VL sequences only. We used OAS updated in 2018[30]. The OAS has rich metadata such as species, vaccinated history, and diseases. We use species to annotate species tag for CTRL method. Because cancers are not in OAS glossed diseases, unknown tag([Unk]) is used for whole OAS dataset. Each VH and VL can be divided by seven regions including three CDRs and four frames. OAS already divided regions in IMGT numbering scheme, yet our in-house patent dataset is not. ANARCI[29] is used to separate regions in IMGT scheme for patent dataset. ANARCI also provides prediction about species of given VH and VL sequence thus, ANARCI predicted species is used for species tag of patent dataset. To address fair training, each dataset is randomly split to train, validation and test set as ratio of 6 : 2 : 2 respectively. While training, only training set is used for updating model and a loss on validation set is observed to choose hyperparameters and stop training before the model suffers overfitting. Finally, the model measures its performance on test set.

### 3.2 Language modeling and attention mechanism

Language modeling is a sequential model that allocate probability, *P* (*x*), of a whole sequence *x* with length *l*. *P* (*x*) can be decomposed to conditional probabilities of next word[31, 18] or character[32] at *i*-th position, *x_i_*, for given previous words, or characters, *x_<i_*:

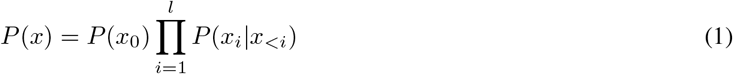

Rather than modeling the probability of a entire sequence, approximating conditional probability of next token(word, character or amino acid) for previous ones gives advantages for model simplicity and sampling efficiency. For example, neural network function *f* (*x_<i_*) can be converted to conditional probability with softmax function:

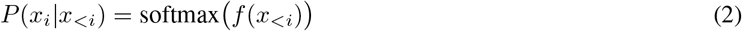

where the softmax function is

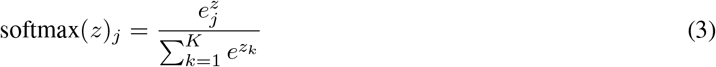

where *K* denotes that number of vocabulary and, *j* and *k* are the indices for tokens in the vocabulary. Many neural network architectures can be constructed to estimate the conditional probability such as linear logistic regression[33, 34], recurrent neural network(RNN)[35], and convolutional neural network(CNN)[36].

As mentioned in section 2, we use GPT-2 architecture. GPT-2 is one of the attention-based neural network model which is now widely used not only in language model[11], but also in computer vision[37]. Attention mechanism actively calculates relationships between a word and the others, in self-attention, including itself. First, scaled dot-product self-attention function[11] maps an input vector to queries, keys and values denoted as *Q*, *K* and *V* respectively. With *Q*, *K*, and *V*, weights between query and keys are calculated and multiplied to values as attention output:

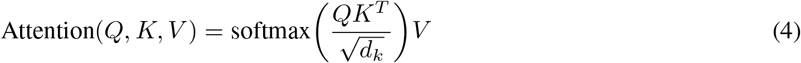

where *d_k_* is a dimension of key vector.

### 3.3 Implementation of GPT-2

GPT-2 is implemented from Huggingface Transformers python package[38] built on pytorch framework[39]. The number of layer, embedding dimensions, hidden layer dimensions and a dropout rate are 12, 252, 1024 and 0.3 respectively. The hyperparameters and early stopping criteria are decided by validation set loss as a result of grid search using Weights and Biases[40].

### 3.4 Materials

Rabbit HRP-conjugated anti HA tag antibody was purchased from Bethyl (USA). Isopropyl-*β*-D-thiogalactoside (IPTG) was obtained from LPS Solution (KOREA). Ni-NTA resin and 1-Step™ Ultra TMB-ELISA Substrate Solution were purchase form Thermo scientific (USA). Disposable column was obtained from BIO-RAD (USA). NC membrane was obtained from GE Healthcare Life Science(Germany). Skim milk was obtained from BD (USA). The Human PD-1 Protein and the human PD-L1 protein were purchased from BIOSYSTEMS Acro (USA).

### 3.5 Selection of antigen binder

AI-designed scFvs were digested with SfiI restriction enzyme (New England Biolabs, USA), and cloned into the pCombi3x vector or pCDisplay-4 vector. 100 ng of recombinant scFv plasmid were incubated with ER2738 bacteria competent cell on ice for 20 min. Subsequently, the competent cells with DNA were incubated at 42 °C for 90 s at heat block, followed by 4 °C incubation for 10 min. LB medium was added to the samples by four times of the competent cell volume, and incubated at 37 °C, 180 rpm for 1 h before being spreaded on LB plate containing carbenicillin in the concentration of 50 *μ*g/mL, and the plates were incubated overnight at 37 °C. To detect whether the scFvs bind to antigen (PD-1 and PD-L1) a number of clones were screened by immunoassay. 750 *μ*L of SB medium (20 g of yeast extract, 30 g of tryptone, 10 g of MOPS, 1 L of pH 7.0) containing carbenicillin was added to each well of the 2.2 mL polypropylene deep well (Axygen, USA). Colonies of the scFv transformants were put in each well, and incubated at 37 °C, 180 rpm for three to four hours. Subsequently, scFv proteins in each well was induced by adding 1mM IPTG at 30 °C, 180 rpm for 20 h. After 20 h induction, the deep well plate was centrifuged at 4000 rpm for 20 min to remove the supernatant and the pellet was resuspended with 400 *μ*L of STE (20 % sucrose, 50 mM Tris-Cl pH 8.0, 1 mM EDTA) buffer. 100 *μ*L of lysozyme (10 mg/mL) was added to the each well and incubated on ice at 180 rpm for 10 min followed by 4 °C incubation at 180 rpm for 10 min with addition of 50 *μ*L 1 M MgCl_2_ to remove outer membrane and peptidoglycan layer. The supernatant containing the periplasmic fraction was obtained after 4000 rpm, 20 min centrifugation to get the soluble scFv proteins. ScFv proteins bound to target antigen were selected by enzyme-linked immunosorbent assay(ELISA). Each 80 *μ*L of the periplasmic extract was treated to target antigen coated 96 well MaxiSorp plate and incubated at 25 °C for 2 h. HRP-conjugated anti-HA antibody (1:3000 diluted), 3,3’,5’5’-tetramethylbenzidine solution, and 2.5 NH_2_SO_4_ were sequentially treated. The clones detected at 450 nm absorbance were selected as target antigen binding antibodies. The expression level of each clone was verified by dot blot. Each 4 *μ*L of the periplasmic extract was treated to NC membrane and incubated at 25 °C for 1h with blocking buffer (skim milk 5 % in PBST(0.05 %)).The blocked NC membrane was incubated with HRP-conjugated anti-HA antibody (1:3000) at 25 °C for 1 h and detected by chemiluminescence.

## 4 Discussion

To overcome limitation of b-cell library in therapeutic antibodies discovery, we suggest library-free active learning method by orchestrating deep learning model and lab experiments. As a proof-of-concept, anti-PD-1 and anti-PD-L1 antibodies are generated, validated, and updated to the training data. Although not all of generated sequences are human nor binding to target, novel antigen bonded sequences are founded by our model with in-house patent dataset. In conclusion, our model shows that deep learning model have potential to discover therapeutic antibody candidates.

